# Minimization of metabolic cost of transport predicts changes in gait mechanics over a range of ankle-foot orthosis stiffnesses in individuals with bilateral plantar flexor weakness

**DOI:** 10.1101/2022.10.14.512205

**Authors:** B. Kiss, N.F.J. Waterval, M.M. van der Krogt, M.A. Brehm, T. Geijtenbeek, J. Harlaar, A. Seth

**Affiliations:** Department of Biomechanical Engineering, Delft University of Technology, Delft, the Netherlands; Amsterdam UMC location University of Amsterdam, Rehabilitation Medicine, Meibergdreef 9, Amsterdam, The Netherlands; Amsterdam UMC location Vrije Universiteit Amsterdam, Rehabilitation Medicine, Boelelaan 1117, Amsterdam, The Netherlands; Amsterdam Movement Sciences, Rehabilitation and Development, Amsterdam, The Netherlands; Department of Orthopaedics, Erasmus Medical Center, Rotterdam, the Netherlands

## Abstract

Neuromuscular disorders often lead to ankle plantar flexor muscle weakness, which impairs ankle push-off power and forward propulsion during gait. To improve walking speed and reduce metabolic cost of transport (mCoT), patients with plantar flexor weakness are provided dorsal-leaf spring ankle-foot orthoses (AFOs). The mCoT during gait depends on the AFO stiffness where an optimal AFO stiffness exists that minimizes mCoT. The biomechanics of why and how there exists a unique optimal stiffness for individuals with plantar flexor weakness are not well understood. To help understand why, we hypothesized that gait adaptations can be predicted by mCoT minimization. To explain how, we hypothesized that the AFO would reduce the required support moment and, hence, metabolic costs from the ankle plantar flexor and knee extensor muscles during stance and reduce hip flexor metabolic cost to initiate swing.

To test these hypotheses, we generated neuromusculoskeletal simulations to represent gait of an individual with bilateral plantar flexor weakness wearing an AFO with varying stiffness. Predictions were predicated on the goal of minimizing mCoT at each stiffness level, and the motor patterns were determined via dynamic optimization. The simulation results were compared to experimental data from subjects with bilateral plantar flexor weakness walking with varying AFO-stiffness.

Our simulations demonstrated that minimization of mCoT predicts gait adaptations in response to varying AFO stiffness levels in individuals with bilateral plantar flexor weakness. Initial reductions in mCoT with increasing stiffness were attributed to reductions in quadriceps metabolic cost during midstance. Increases in mCoT above optimum stiffness were attributed to the increasing metabolic cost of both hip flexor and hamstrings muscles.

The insights gained from our simulations could inform clinicians on the prescription of personalized AFOs. With further model individualization, simulations based on mCoT minimization may sufficiently predict adaptations to an AFO in individuals with plantar flexor weakness.

**Author Summary:** Neuromuscular disorders like stroke, Charcot-Marie-Tooth disease, and poliomyelitis often lead to calf muscle weakness, which makes walking slower and more demanding. To improve walking speed and reduce energy demand, patients with calf muscle weakness are frequently provided ankle-foot orthoses (AFOs). The energy demand of walking is affected by the AFO’s stiffness and there is a stiffness that minimizes the energy demand for an individual with calf weakness. To uncover the optimal stiffness, we generated simulations of an individual with calf muscle weakness walking with an AFO over a range of stiffnesses. Stable walking patterns were generated that minimized the energy demand for a given stiffness. We found that the initial reductions in energy demand as stiffness increased, were attributed to reductions in quadriceps muscle energy. Increases in energy demand as stiffness increased above the optimum were attributed to the increased energetic cost of both hip flexor and hamstrings muscles. With further model individualization, we believe that simulations based on minimizing the energy demand of movement can sufficiently predict adaptations to an AFO. Simulations can enable the prescription of personalized AFOs for individuals with neuromuscular disorders that help them walk with sufficient speed and efficiency to keep up with their peers.

## Introduction

The plantar flexor muscles, consisting of soleus and the gastrocnemius, are often weakened in persons with neuromuscular disorders, such as Charcot-Marie-Tooth disease and poliomyelitis [1][2]. Weakness of the plantar flexors results in an altered gait pattern, characterized by reduced push-off power, and excessive ankle dorsiflexion and knee flexion during stance [3][4]. These gait deviations lead to a lower walking speed [5] and an elevated metabolic cost of transport (mCoT) [6], which limits daily physical mobility [7]. Dorsal leaf spring (DLS) ankle-foot orthoses (AFOs) are often prescribed to provide mechanical support during stance, augment push-off, and hence to reduce mCoT. In a DLS-AFO, a leaf spring connects a footplate to a calf casing posterior of the ankle and passively restricts ankle dorsiflexion by producing an external plantarflexion moment when the ankle is dorsiflexed. As a spring, the AFO can store energy when moving into dorsiflexion and release this energy as the ankle moves towards plantarflexion, thereby providing additional positive work during push-off [8].

In individuals with plantar flexor weakness, the effects of an AFO on improving gait kinematics and kinetics and reducing mCoT have been shown to depend on the stiffness of the leaf spring [9][10][11]. Beginning at low and with increasing AFO stiffnesses, the mCoT first decreases, before increasing at higher stiffness levels, demonstrating a convex relation between AFO stiffness and mCoT with an optimum stiffness where mCoT is minimal [9][10]. As demonstrated in healthy individuals [12][13][14], minimizing mCoT is prioritized during gait. As such, it can be expected that patients with gait disorders prefer walking with the stiffness that minimizes their metabolic energy cost. In case of plantar flexor weakness, the initial reduction in mCoT is thought to be the result of normalizing ankle and knee angles and moments which requires adequate AFO stiffness [9][10]. Normalization of the ankle and knee biomechanics is hypothesized to lead to a decrease in the metabolic cost of the quadriceps muscles and thereby reduce mCoT [15]. The initial decrease in mCoT may be further explained by a reduction in the metabolic cost of the plantar flexors as the AFO replaces the biological ankle plantarflexion moment during stance [11][16][17]. However, at higher stiffnesses, as the AFO restricts the ankle range of motion (RoM) [9][18][16], it limits active biological ankle power generation and energy storage and release [19][16] of the AFO during push-off [9][10]. The reduced ankle push-off work may result in higher energy losses at contralateral heel-strike and lead to compensatory hip flexion work to initiate the swing phase [16], which are potential causes for the increased mCoT at higher AFO stiffness levels. However, how each of these factors contribute to the relation between AFO stiffness and mCoT in people with plantar flexor weakness is unknown.

The aim of this study was to gain insights into why and how mCoT is affected by AFO stiffness variation in individuals with plantar flexor weakness by using predictive musculoskeletal simulations. First, based on the assumption that mCoT is a predictor for gait pattern changes in healthy individuals [14][12], we hypothesized that minimization of mCoT is a predictor of kinematic, kinetic, and mCoT changes to varying AFO stiffness in individuals with bilateral plantar flexor weakness. Second, we tested whether initial reductions in mCoT with increasing stiffness are explained by i) decreasing metabolic cost of the quadriceps as the knee moments are normalized, and ii) decreasing metabolic cost of the plantar flexors as the AFO replaces the ankle plantar-flexion moment during stance phase. Third, we hypothesized that increases in mCoT as stiffnesses exceed the optimum stiffness are caused by the increasing metabolic cost of hip flexor muscles to initiate the swing phase as total push-off power decreases.

## Methods

We created a planar musculoskeletal model of an individual with bilateral plantar flexor weakness, similar to [20] using OpenSim [21][22], and implemented an AFO with varying stiffness. To generate predictive gait simulations, we employed a reflex-based neuromuscular controller and optimized the control parameters using dynamic optimization to minimize mCoT, and solved the optimization problem in SCONE [23][24][20]. Predictive simulation results were compared to experimental data of subjects with bilateral plantar flexor weakness walking with varying AFO stiffness [10].

### Musculoskeletal model

Based on the model of Delp et al. [25][24], we created a model with 9 degrees of freedom (3 around the pelvis and 1 around the hip, knee and ankle of each leg), actuated by 9 Hill-type muscles per leg (tibialis anterior, soleus, gastrocnemius, vasti, rectus femoris, biceps femoris short head, biarticular hamstrings, iliopsoas, gluteus maximus) in OpenSim 3.3 [21][22]. We set the maximum isometric muscle strength of the soleus and gastrocnemius muscles of both legs to 40% of normal healthy values (i.e. 60% muscle weakness), to induce bilateral plantar flexor weakness. Additionally, we restricted the ability to activate the plantar flexor muscles to 50%, to take into account that the weakened muscles would completely fatigue if they would be maximally activated for 10-20% of the gait cycle [26][27][28]. We modified passive muscle and tendon parameters in the model to maintain similar passive muscle forces as in the healthy model [20][29]. We set the slow twitch fiber ratios according to Johnson et al.[30] and Garrett et al.[31], similarly to Ong et al.[24]. We scaled the model according to experimental marker data of one subject with bilateral plantar flexor weakness close to the group’s mean height (177 cm) and body mass (81 kg) from the experimental study [10].

To model the forces between the ground and the foot, we used a compliant Hunt-Crossley contact model [32]. We placed one contact sphere at the heel and one at the toe of each foot (Fig 1). We set the force parameters (stiffness, dissipation and friction) according to Veerkamp et al.[33], and modelled the knee ligaments using a rotational spring (2 Nm/deg) and damper (0.2 Nm/deg/s) around the knee joint if the knee angles were outside the 5-120 deg flexion range [20].

**Fig 1.**
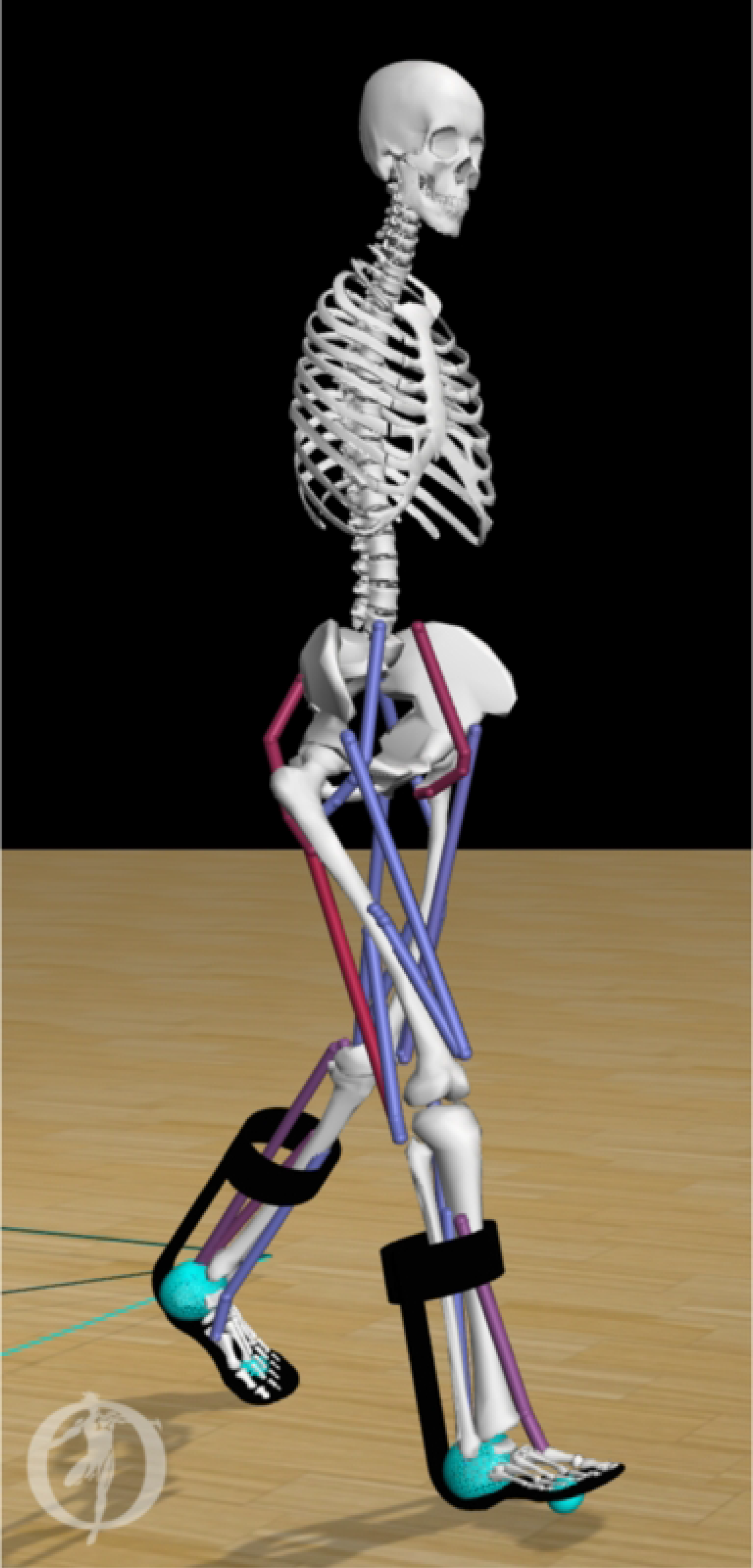
Musculoskeletal model of an individual with bilateral plantar flexor weakness equipped with AFOs. The stiffness (range 0 − 7 Nm/deg) of the dorsal leaf spring is modelled in OpenSim as a torsional spring between the calf-casing and footplate of each AFO (black). Contact between the foot and the ground are modelled by forces generated by compliant contact spheres (cyan).

We modelled each AFO as two rigid parts, including a calf casing and a footplate with their experimental mass (calf casing: 0.2 kg, footplate and shoe: 0.5 kg) [10]. We attached the AFO parts rigidly to the tibia and calcaneus, respectively (Fig 1). We modelled the stiffness of the AFO as two linear torsional springs for ankle dorsiflexion and for plantarflexion. In order to match experimental movement of the ankle within the AFO, the AFO did not deliver a moment in the neutral angle range, i.e. between 4.5 deg plantarflexion and 2 deg dorsiflexion. In DLS-AFOs, this small range depends on the material and manufactured geometry of the AFO, and its fit on the subject’s leg. The neutral angle range was defined from the ankle angle range during swing phase of the subject because in swing phase the AFO exerts only small moments on the ankle joint [34].

### Simulation framework

We used SCONE (v1.6.0), a simulation, control, and dynamic optimization framework [23], to simulate gait of 10 seconds in duration. The muscle activations were generated by a reflex-based gait controller [35][24], whose parameters were optimized by minimizing the specified objective function using the Covariance Matrix Adaptation evolutionary strategy (CMA-ES) [36][20][24][37].

The objective function (J) was comprised of desired high-level tasks during gait, where minimization of mCoT (*j*_mCoT_) was the primary measure. The following measures were included in the objective function to be minimized:

*J*_mCoT_ was the mCoT measure, which aggregated the total muscle metabolic cost divided by the distance travelled. We computed the metabolic cost of each individual muscle, according to the muscle metabolic model by Uchida et al. [38].

P_Gait_ was a penalty on the deviations of the simulation from gait at a specified minimum velocity of 1.22 m/s, to match experimental data without falling down [10].

We added P_DOFLimAnkle_ and P_DOFLimKnee_ penalties to keep the ankle angle and passive knee forces within physiological limits. We gave penalties, when the ankle angle was outside of the [−60, 60] deg range and when the absolute coordinate limit moment acting on the knee joint was larger than 5 Nm [20].

P_FGImpact_ was a penalty composed of the sum of the absolute ground reaction force derivative over the simulation divided by the distance travelled, which was included to penalize high loading rates at impact.

P_HeadStab_ was a penalty for excessive head accelerations calculated as the sum of the absolute head accelerations normalized by distance travelled [39][40].

The weights associated to these high-level tasks were w_mCoT_=1.5, w_Gait_=10^9^, w_DOFLimAnkle_=0.1, w_DOFLimKnee_=0.01, w_FGImpact_=0.05, w_HeadStab_=0.1. We chose the weights based on a previous study [20], but adapted with a higher emphasis on J_mCoT_ to test the hypothesis that energy cost minimization can predict gait changes with AFO stiffness.

We ran the simulations for stiffness levels between 0 and 7 Nm/degree, with steps of 1 Nm/degree. We ran 5 optimizations in parallel with different random seeds in each round. Each optimization was terminated when the average reduction of the cost function score in the last 200 generations was smaller than 0.01 %. As initial guess, we used a controller with parameters resulting in healthy gait [20]. We set the initial step size (*σ*), as the initial standard deviation of the parameters, to 0.05 [41].

To check the robustness of our results, we ran multiple optimizations in sequence. We used the best results of an optimization to initialize the next optimization with the same model (same AFO stiffness setting) and the same initial step size, similar to Song et al. [37] and Ong et al. [41]. Since the trend of the results was not changing qualitatively between the first and second round of optimizations, we performed only two rounds of optimizations. We considered the results of the second round as the final results.

### Comparison with experimental data

To test whether mCoT is a good predictor of the gait changes with varying AFO stiffness, we compared our predictive simulations with experimental gait and mCoT data of 24 bilateral weakness subjects walking with 5 different AFO stiffness configurations (2.8, 3.5, 4.3, 5.3, 6.6 Nm/deg) [10]. In this experimental study, the mean mCoT (in J/kg/m) was evaluated from a 6-min comfortable walk test with simultaneous gas-analysis over the last three minutes of the test. The study assessed gait kinematics, kinetics and ground reaction forces with a 3D gait analysis on a 10-meter walkway using the PlugInGait marker model [42]. Based on these measurements, clinically important gait features for the evaluation of AFOs, e.g. the peak ankle dorsiflexion angle, ankle power, knee angle, knee moment and AFO-generated power, were calculated using a custom-made script in MATLAB® R2015b (MathWorks Inc.).

To calculate the joint moments from the predictive simulations, we processed the simulation results with the Analysis Tool in OpenSim. Based on the joint angles and moments, we calculated the joint powers (Fig 2). To calculate negative and positive joint work over the whole gait cycle and separate gait phases we used trapezoidal numerical integration of the joint power. We divided the gait cycle into gait phases according to the definitions of Whittle [43]. These gait phases were: loading response, midstance, push-off comprising of terminal stance and preswing, and swing. In order to assess the source of the mCoT in detail, we calculated the simulated muscle metabolic energy cost [38] over whole gait cycles, and different gait phases, for all 9 muscles. We normalized the metabolic cost by body mass, mean walking speed and simulation duration to get the total metabolic cost over a gait cycle in J/kg/m. We used a custom-made script in MATLAB® R2020b (MathWorks Inc.) for all calculations.

**Fig 2.**
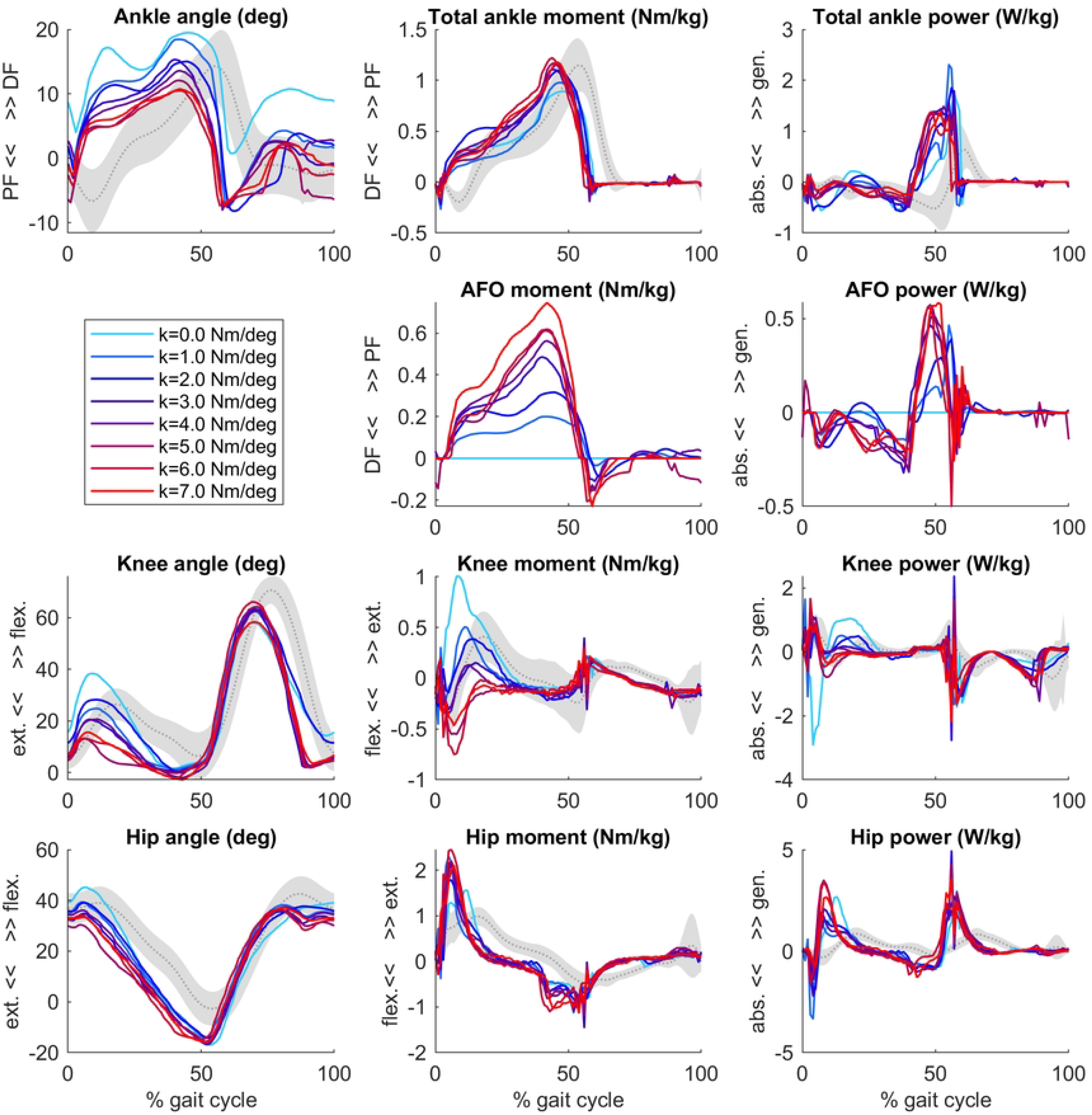
Ankle, AFO, knee and hip angles, moments and powers from the forward simulations. Simulation data of a typical patient model with bilateral calf muscle weakness wearing an AFO with varying stiffness levels (range 0 − 7 Nm/deg, coloured lines). Grey curves and shading represents patient data without AFO (mean ±1 SD) [10]. After initial contact, the ankle, knee and hip joints became more extended as AFO stiffness increased.

To compare simulation-based outcome measures to experimental observations, we calculated the effect of 1 additional Nm/degree in stiffness for both the simulations and experimental data using a linear fit across the stiffness levels for the following key gait features: peak ankle dorsiflexion angles, peak total-, biological- and AFO ankle joint moments and powers, peak knee extension angles and peak knee joint moments (between 35-50% of gait cycle) [10]. We assessed the goodness of fit of the curve by its coefficient of determination value (Rsq), calculated in MATLAB® R2020b (MathWorks Inc.) [44]. To assess the similarity between the simulated and experimentally obtained slopes, we expressed the difference in slope in standard deviation of the experimental slope based on the 24 patients.

## Results

Our predictive gait simulations took 20.46 hours on average to complete on an AMD Ryzen 9 3950X (16 CPUs – 32 virtual cores with hyperthreading, 3.5 GHz base) computer on 10 parallel threads. The primary objective of mCoT minimization contributed to about 90% of the final optimization scores in all simulations. In the final simulations, P_Gait_ and P_DoFLimAnkle_ were optimized to zero and did not contribute to the objective score. An overview of the simulated joint angles, moments and powers for varying AFO stiffness levels are presented in Fig 2.

### Comparison of simulated and experimental results

The predicted slopes of peak total, biological and AFO-provided ankle joint moment and power, peak ankle dorsiflexion angle, peak knee extension angle and peak internal knee flexion moment were all within 1.2 standard deviations (SD) of the experimental data. The highest slope difference was found for peak total ankle moment and peak AFO moment, where a larger effect of additional stiffness was predicted by the simulations than found in experimental data. Peak total ankle moment was approximately constant in the experiments but showed an increasing trend in the simulations (Fig 2, Fig 3 and S1 Table).

**Fig 3.**
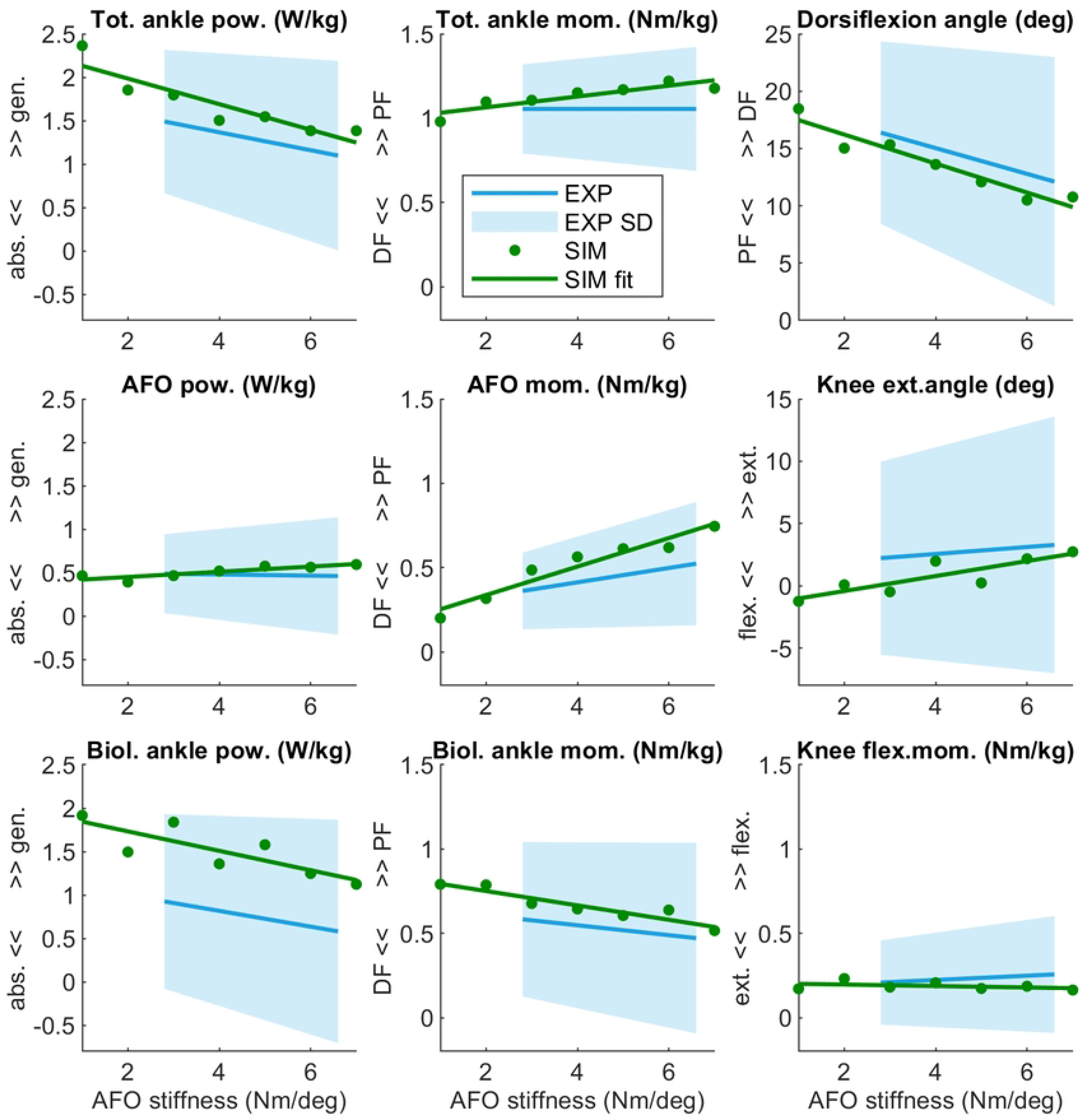
Peaks of relevant joint and AFO powers, moments and angles during the stance phase. AFO stiffness was varied from 1 − 7 Nm/deg. The results of simulation with fitted lines are shown in green, and mean experimental outcomes of all patients in blue. As experimental data the mean and SD (blue shading) of the individual fitted lines were used.

### Simulated AFO effects on mCoT and muscle metabolic consumption

The mCoT showed a clear minimum with increasing stiffness in both the simulated and individual experimental results (Fig 4). The simulations presented a strong quadratic trend, Rsq = 0.836, while the average experimental results showed a less pronounced quadratic trend, Rsq = 0.634, due to large inter-subject variability (Fig 4).

**Fig 4.**
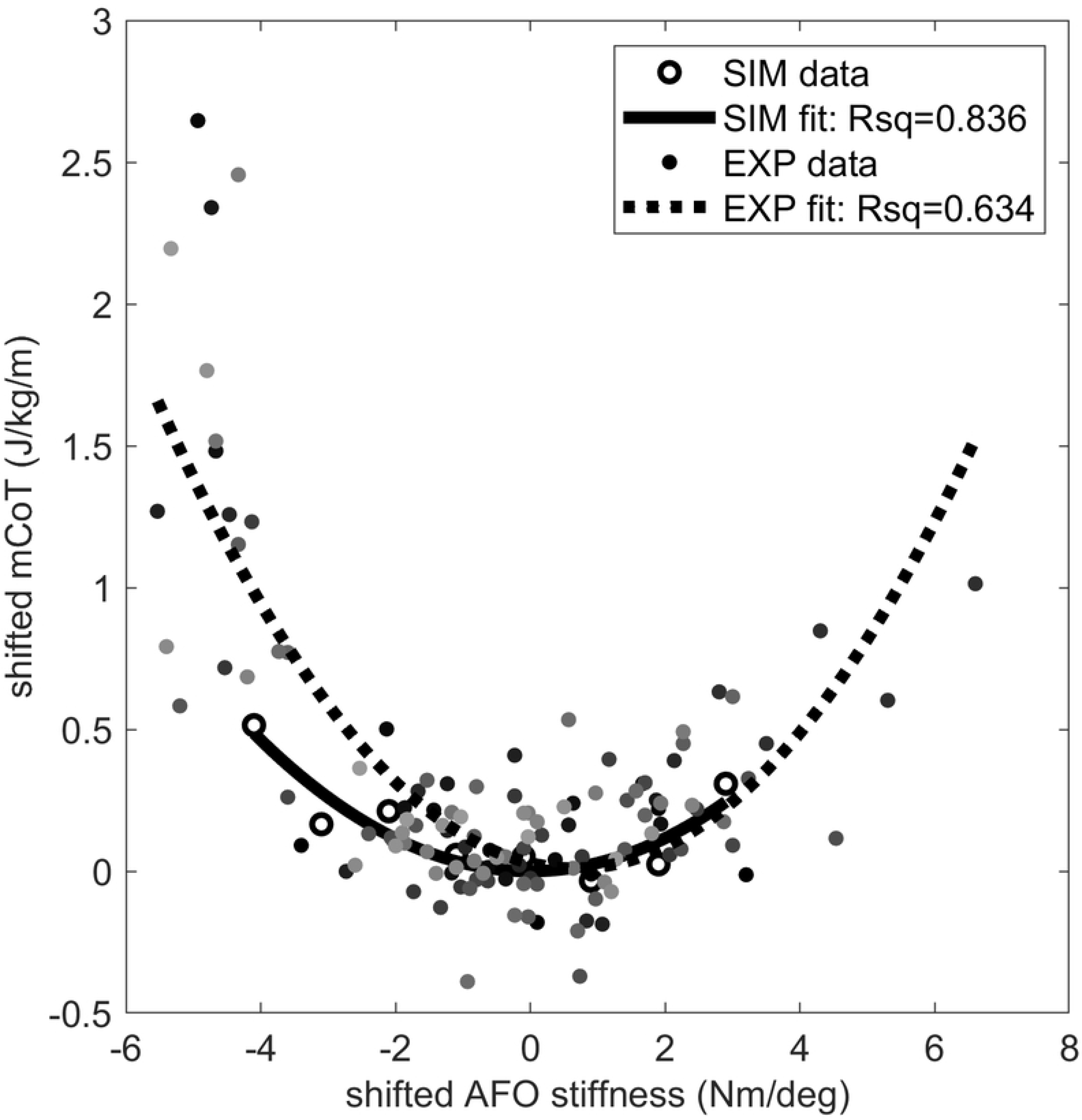
The mCoT in simulation and individual experiments wearing an AFO with varying stiffness. The individual experiments included the condition without an AFO and wearing shoes for 21 subjects. A quadratic curve was fitted to the data points for each subject dataset and the minimum of the fitted curve was taken for each subject as their individual minimum mCoT value which is happening at their individual optimal stiffness. The mCoT values of the subjects were shifted by their minimum mCoT value and the AFO stiffness values were shifted by the subject’s optimal stiffness value. The shifted individual experimental subject data is shown in different shades of grey. One quadratic curve was fitted to all shifted experimental subject data (dotted line), the goodness of fit is indicated by the Rsq number on the plot (Rsq = 0.634). The same was done for the simulation results (solid line, Rsq = 0.836). Both the experimental (EXP – shaded dots) and simulation data (SIM – open circles) show quadratic trends (Rsq > 0.63).

The largest change in metabolic cost of individual muscles was found in the vasti, which also showed a quadratic trend (Rsq = 0.928). In contrast, the metabolic cost of the hamstrings and iliopsoas increased continuously. Both the soleus and gastrocnemius metabolic cost did not change substantially with increasing AFO stiffness (Fig 5 and S2 Table). Gluteus maximus, tibialis anterior, rectus femoris and biceps femoris short head muscles did not show any change with increasing AFO stiffness (S2 Table).

**Fig 5.**
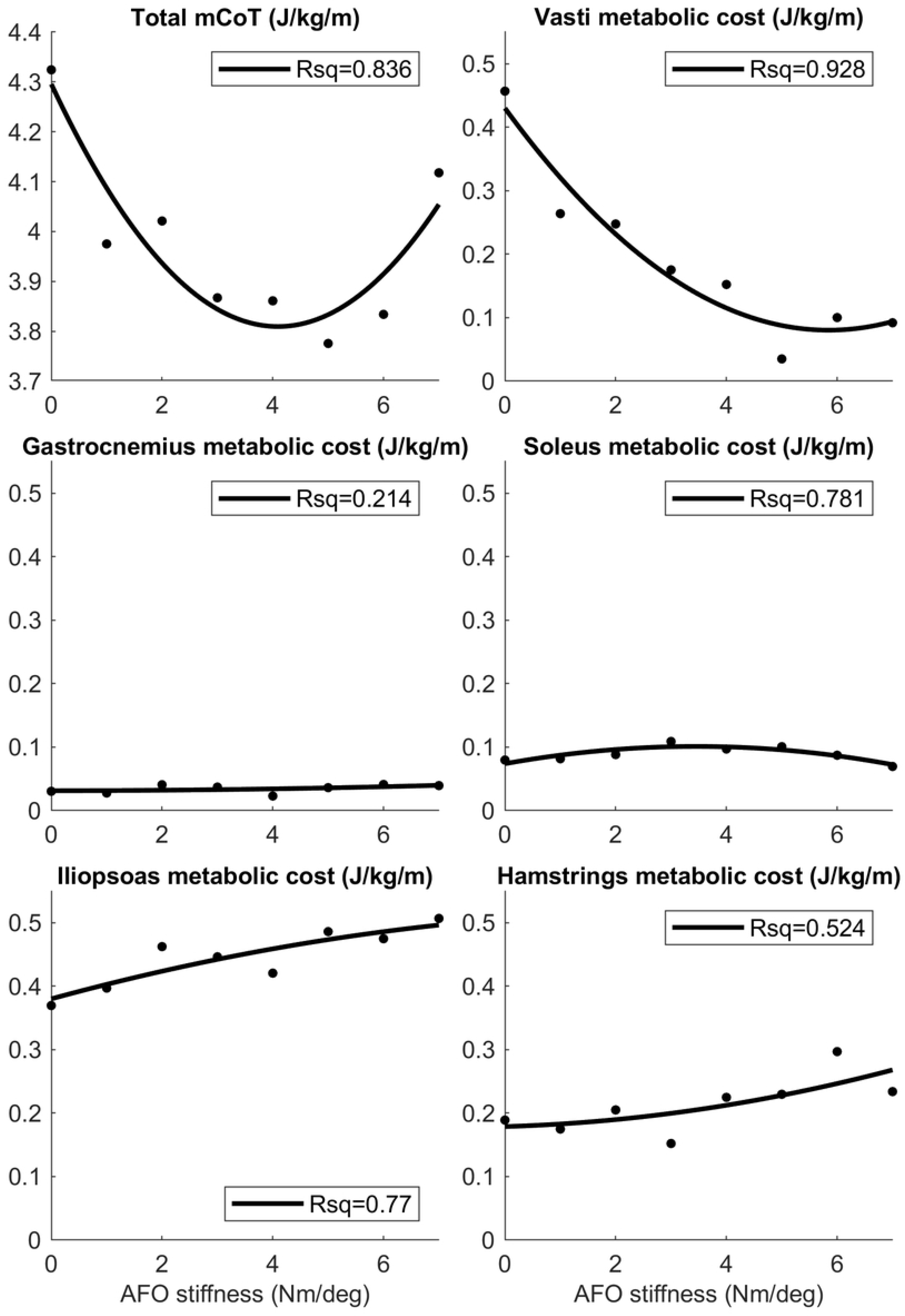
The mCoT, and total metabolic cost of the top 5 muscle contributors in simulations. Total metabolic cost was taken during one whole gait cycle as AFO stiffness was varied from 0 − 7 Nm/deg. Quadratic curves were fitted to the data-points and the Rsq values represent the goodness-of-fit of the curves.

### Joint work and muscle metabolic consumption per gait phase

Total knee joint work did not differ with increasing AFO stiffness during the loading response, while vasti metabolic cost decreased and hamstrings metabolic cost increased, especially above 3 Nm/degree. In midstance, positive knee joint work decreased, negative knee joint work increased and vasti metabolic cost decreased, while hamstrings metabolic cost did not show a clear trend with increasing AFO stiffness. (Fig 6, S3 and S4 Table).

**Fig 6.**
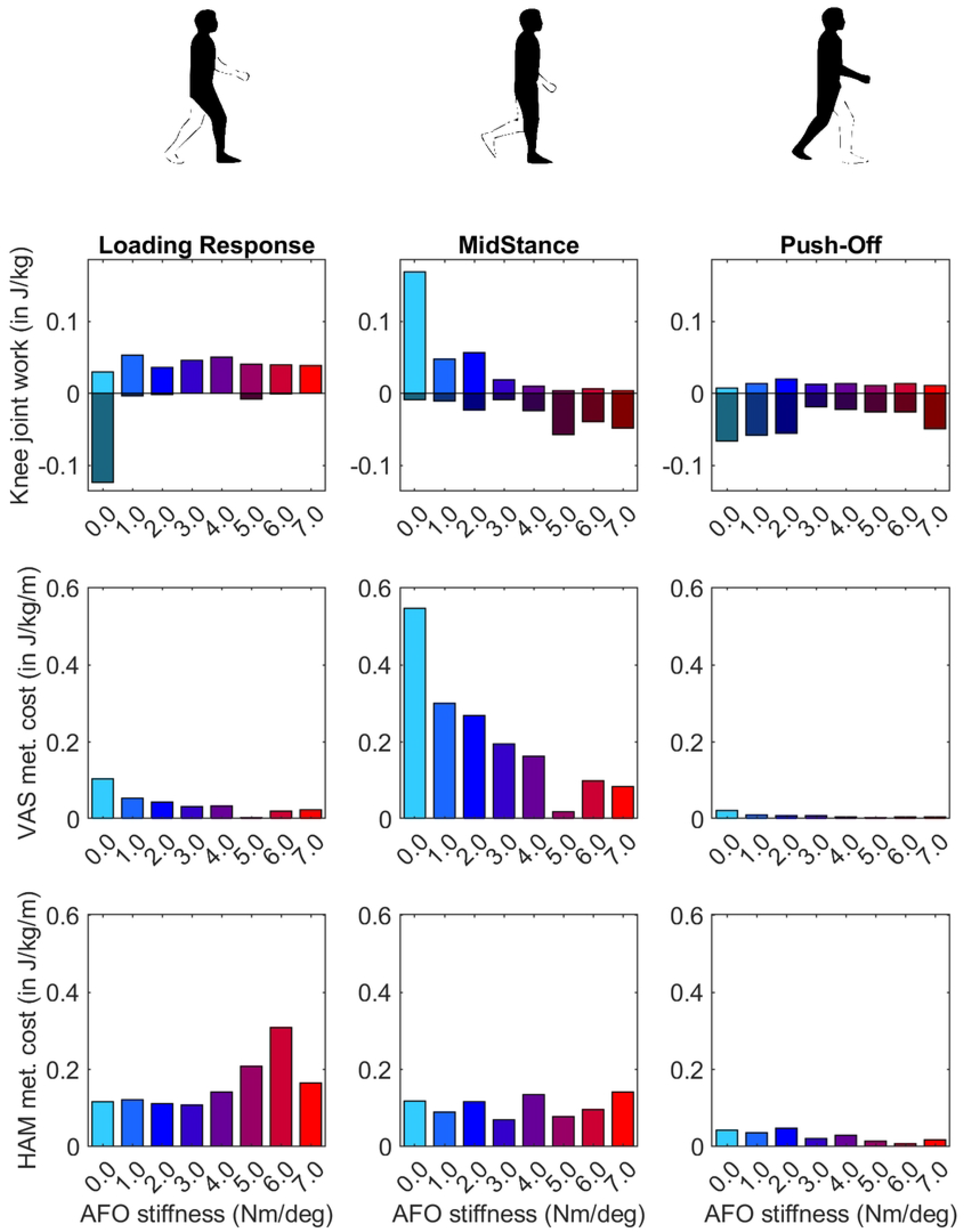
Knee joint work, vasti and hamstrings metabolic cost during loading response, midstance, and push-off. AFO stiffness was varied from 0-7 Nm/deg. Knee joint work was portioned into positive and negative work and these were totalled and reported for each phase of gait.R

During loading response, negative biological ankle work decreased with increasing AFO stiffness, while no effect of stiffness on AFO work was found. Similarly, no effect on the metabolic cost of the soleus or gastrocnemius was found with increasing stiffness. In midstance, negative biological ankle joint work decreased and negative AFO work increased with increasing stiffness. Soleus metabolic cost increased slightly until 5 Nm/deg AFO stiffness. During push-off, biological ankle work, AFO work, and soleus metabolic cost increased until 3 Nm/deg. At higher stiffnesses, biological ankle work generation and soleus metabolic cost decreased again (Fig 7, S3 and S4 Table).

**Fig 7.**
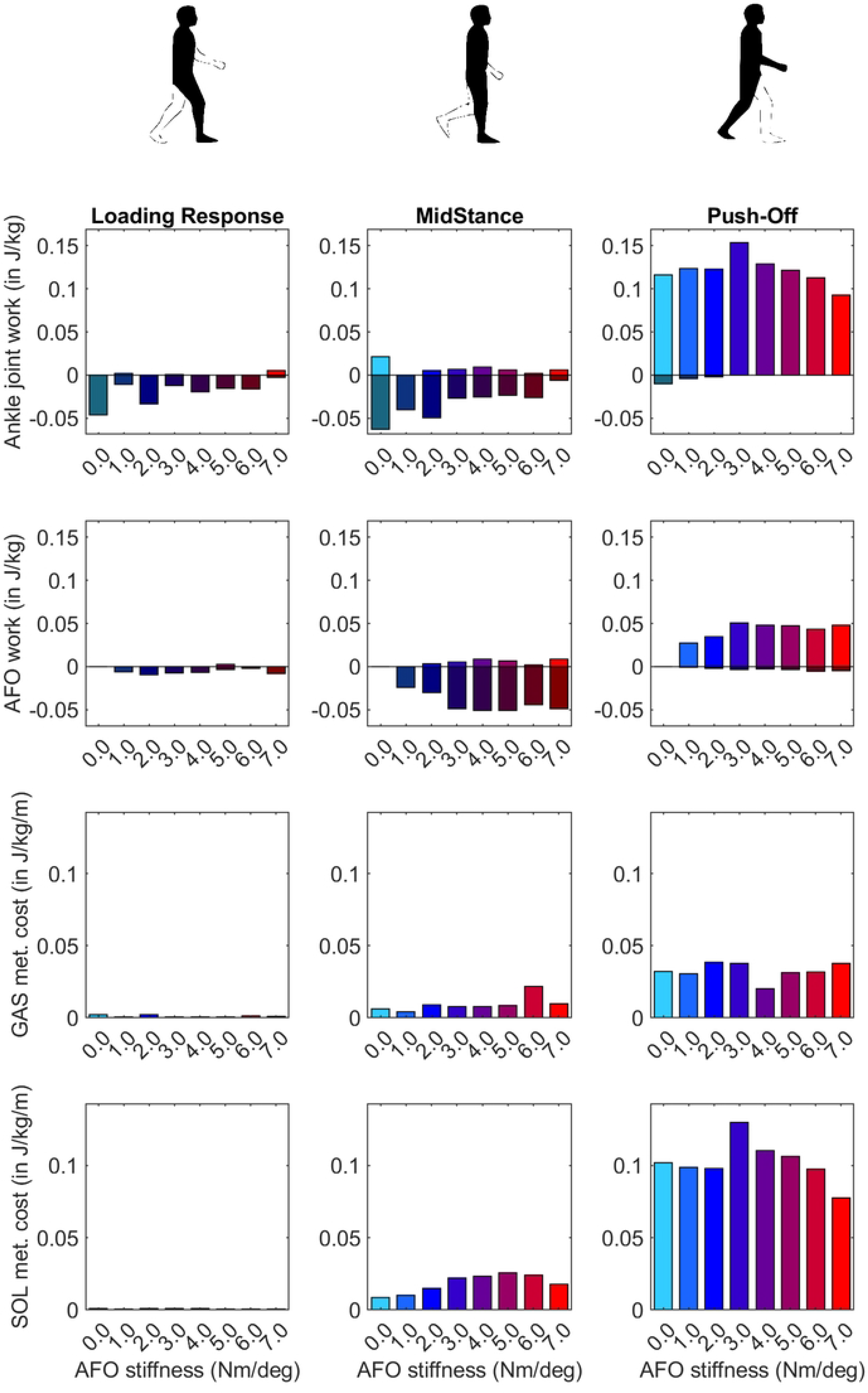
Biological ankle and AFO work, gastrocnemius and soleus metabolic cost during loading response, midstance, and push-off. AFO stiffness was varied from 0-7 Nm/deg. Biological ankle joint and AFO work was portioned into positive and negative work and these were totalled and reported for each phase of gait.

During loading response, negative hip joint work decreased and hamstrings metabolic cost increased with increasing stiffness. In early midstance, positive hip joint work increased, and in late midstance, negative hip joint work and iliopsoas metabolic cost increased with increasing AFO stiffness (Fig 8, S3 and S4 Table).

**Fig 8.**
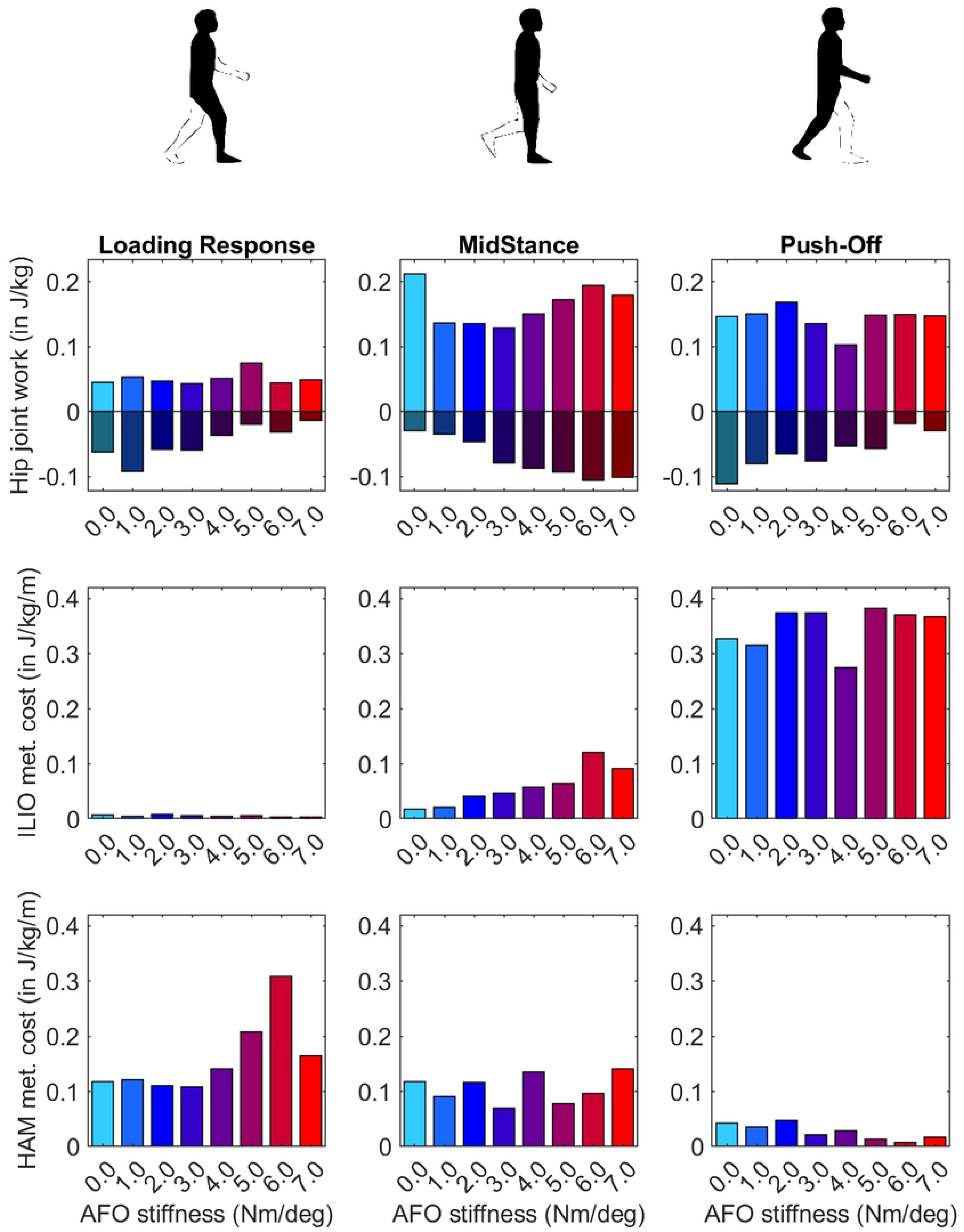
Hip joint work, iliopsoas and hamstrings metabolic cost during loading response, midstance, and push-off. AFO stiffness is varied from 0-7 Nm/deg. Hip joint work was portioned into positive and negative work and these were totalled and reported for each phase of gait.

## Discussion

The aim of this study was to gain insights into why and how motor pattern adaptations in people with bilateral plantar flexor weakness result in an optimal AFO stiffness to minimize mCoT. As hypothesized, our results showed that minimization of mCoT predicts most of the kinematic, kinetic, and mCoT changes due to varying AFO stiffness in this population. Initial reductions in mCoT with increasing stiffness were attributed to reductions in quadriceps metabolic cost, as hypothesized, but in contrast to our hypothesis, plantar flexor metabolic cost did not decrease. Increases in mCoT above the optimum AFO stiffness were attributed to the increasing metabolic cost of both hip flexor muscles and hamstrings muscles.

The mCoT appears to be a good predictor of gait changes in individuals with bilateral plantar flexor weakness wearing AFOs with varying stiffness. Our forward simulations predicted changes in lower extremity kinematics and kinetics due to AFO stiffness variations within 1.2 SD of the experimentally observed changes. Although mCoT was not the only factor within our objective function, it had by far the largest contribution as it was responsible for 90% of the final objective function values. These findings support the claims that humans tend to minimize mCoT when walking as previously demonstrated in healthy subjects [12][13][14] and in simulations of patients walking without assistive devices [45].

Differences between simulated and experimental data were found in the knee joint angle and moment curves (Fig 2). Although the effects of varying AFO stiffness on specific gait features at midstance were predicted reasonably well, in early stance the knee angle and moment became more extended with increasing AFO stiffness (Fig 2 and 3). This effect is likely explained by the fact that more extended knees in early stance are linked to decreased mCoT [46][47][48]. The human likely minimizes or is subjected to other factors, such as muscle fatigue and loading rate [49][50][33], which may cause the knee flexion in loading response instead of the metabolically more efficient straight knee. Although loading rates were penalized in our simulations, loading rates still increased up to twice as much as found in healthy subjects as AFO stiffness increased (S1 Fig), which could increase the risk of lower limb stress fractures [51][52].

The convex mCoT trend with respect to AFO stiffness can be explained by the metabolic cost changes in the quadriceps (vasti), hip flexor (iliopsoas) and hamstrings muscles. This parabolic trend was also present in the individual experimental data [10] (Fig 4). As hypothesized, initial reductions in mCoT starting at low and with increasing AFO stiffness were due to a decrease in the metabolic cost of the quadriceps (vasti) muscles (Fig 5). From low to medium AFO stiffnesses, the knee angle and moment normalized, reducing the metabolic cost of the vasti [46][48]. At higher stiffness levels, the knee became increasingly extended, which minimized mCoT but might cause knee pain in real life [51][52]. Contradicting our hypothesis and experimental data in healthy subjects [53], metabolic cost of the plantar flexor muscles did not decrease with increasing AFO stiffness. As muscle activation and metabolic cost changes are related factors, our simulation result was also contrary to the findings of Harper et al. [18] who found reductions in medial gastrocnemius muscle activation with increasing AFO stiffness in lower limb salvage patients [18]. We likely did not observe reductions in plantar flexor metabolic cost, because the low muscle strength in the model resulted in proportionally low muscle mass, which reduces the muscle’s contribution to metabolic cost [29], and hence, even without an AFO the plantar flexors did not contribute substantially to mCoT.

In agreement with our hypothesis, increases in mCoT above the optimum stiffness were partly due to increases in the metabolic cost of the hip flexor (iliopsoas) muscles. Iliopsoas metabolic cost increased at the end of midstance before the start of push-off, potentially as a pre-activation to help initiate the swing phase. Increased hip work was also shown in an experimental study in patients with chronic stroke or multiple sclerosis where 0.5-5.4 Nm/deg AFO stiffnesses were tested [16]. Additionally, an increase in metabolic cost of the hamstrings muscles added to the increase in mCoT above the optimum stiffness. This metabolic cost increase can be seen during early stance where slightly more extended hip joint angles, larger hip extension moments (Fig 2) and decreasing negative hip joint work (Fig 8) can also be observed as AFO stiffness increases. At high stiffness levels, the hip is more extended at initial contact, and hip flexion is reduced during loading response that further contributes to the increased knee joint loading rates at high stiffness.

### Limitations and future work

To test our hypotheses, we used a simplified, planar musculoskeletal model where medio-lateral stabilization was excluded, which could explain why our mCoT results were ∼10% lower [54] than in the experiments. With suboptimal AFO settings, the out-of-plane compensation, such as trunk motions, are known to be more extreme [55], hence the sensitivity of the mCoT trend to AFO stiffness could be higher in reality than in our simulations.

In the experimental study that was used for comparison, only AFOs with a stiffness in the range of 2.8-6.6 Nm/deg were tested. Hence, we were unable to verify the validity of our prediction with low stiffness levels.

In the future, simulations might be used to predict the individual optimal AFO properties, for which improving our understanding of the underlying mechanisms in this study was an essential first step. To be able to accurately predict adaptations to an AFO at the level of an individual, out-of-plane degrees of freedom and muscle actions should be investigated to understand the affects out-of-plane compensations. Furthermore, sensitivity analyses should be performed to evaluate the effect of patient and device characteristics, such as body weight, muscle weakness and muscle spasticity, and the neutral angle range of the AFO, have on the optimal AFO stiffness. With individualized models, our forward simulations could help predict the individual adaptations of patients to an AFO and improve the prescription of AFO settings.

## Conclusion

We showed that adaptations in gait mechanics due to varying AFO stiffness, in individuals with bilateral plantar flexor weakness, can be predicted by minimization of mCoT. Our simulation results demonstrate the convex relation between mCoT and AFO stiffness, and are able to explain this shape by decreases in quadriceps metabolic cost in midstance, and increases in metabolic cost of the hamstrings during loading response and iliopsoas in mid-to-late stance as AFO stiffness increases above the optimal. In the future, mCoT minimization may enable predictions for individualized gait adaptations to an AFO for people with bilateral plantar flexor weakness and facilitate optimal AFO prescriptions. The musculoskeletal models (in OpenSim, https://simtk.org/projects/opensim) and code, which were used to execute the gait optimization (in SCONE, https://simtk.org/projects/scone), and our complete simulation results are provided at https://simtk.org/projects/afo-predictions.

## Acknowledgments

This project has been made possible in part by grant 2020-218896-5022 (AS) from the Chan Zuckerberg Initiative Donor-Advised Fund (DAF), an advised fund of Silicon Valley Community Foundation.

This project was supported by the Netherlands Organization for Health Research and Development (ZonMw) IMDI grant 104022003 (NFJW).

## Supporting information

**S1 Fig. Peak loading rate (GRF derivative) in loading response across 0-7 Nm/deg stiffnesses (BW: body weight).** The found peak loading rate values are more than twice as high as the loading rate on healthy subjects during walking at 1.3 m/s [51], but lower than the loading rate on healthy subjects during running at 3.7 m/s [56].

**S1 Table. Slope of relevant joint kinematics, kinetics and powers during the stance phase of gait.** AFO stiffness was varied from 1 − 7 Nm/deg. From left to right: slope of the line fitted to the forward simulation result, mean of the slope of the lines fitted to the bilaterally affected patients’ results, standard deviation (SD) of the slope of the lines fitted to the bilaterally affected patients’ result, difference between the simulation slopes and the mean of the experimental slopes divided by the standard deviation of the experimental slopes.

**S2 Table. Mean metabolic cost for the whole model and for each muscle group of the model.** Data taken during a whole gait cycle as AFO stiffness was varied from 0 − 7 Nm/deg.

**S3 Table. Positive and negative mechanical joint work calculated with integration for each gait phase.** AFO stiffness was varied from 0 − 7 Nm/deg.

**S4 Table. Metabolic cost of transport of each muscle group calculated with integration for each gait phase.** AFO stiffness was varied from 0 − 7 Nm/deg.

## References

1. Rossor AM, Murphy S, Reilly MM. Knee bobbing in Charcot-Marie-Tooth disease. Pract Neurol [Internet]. 2012 Jun [cited 2022 Jun 21];12(3):182–3. Available from: https://pmc/articles/PMC3736802/

2. Neumann DA. Polio: Its impact on the people of the United States and the emerging profession of physical therapy [Internet]. Vol. 34, Journal of Orthopaedic and Sports Physical Therapy. Movement Science Media; 2004 [cited 2022 Jun 21]. p. 479–92. Available from: https://www.jospt.org

3. Steele KM, Rozumalski A, Schwartz MH. Muscle synergies and complexity of neuromuscular control during gait in cerebral palsy. Dev Med Child Neurol [Internet]. 2015 Dec 1 [cited 2021 Nov 10];57(12):1176–82. Available from: https://pmc/articles/PMC4683117/

4. Ploeger HE, Bus SA, Nollet F, Brehm MA. Gait patterns in association with underlying impairments in polio survivors with calf muscle weakness. Gait Posture. 2017 Oct 1;58:146–53.

5. Waterval NFJ, Brehm MAA, Ploeger HE, Nollet F, Harlaar J. Compensations in lower limb joint work during walking in response to unilateral calf muscle weakness. Gait Posture [Internet]. 2018 Oct 1 [cited 2021 Feb 3];66:38–44. Available from: https://linkinghub.elsevier.com/retrieve/pii/S0966636218314255

6. Brehm MA, Nollet F, Harlaar J. Energy demands of walking in persons with postpoliomyelitis syndrome: Relationship with muscle strength and reproducibility. Arch Phys Med Rehabil. 2006;87(1):136–40.

7. Nollet F, Beelen A, Prins MH, De Visser M, Sargeant AJ, Lankhorst GJ, et al. Disability and functional assessment in former polio patients with and without postpolio syndrome. Arch Phys Med Rehabil. 1999 Feb 1;80(2):136–43.

8. Hegarty AK, Petrella AJ, Kurz MJ, Silverman AK. Evaluating the effects of ankle-foot orthosis mechanical property assumptions on gait simulation muscle force results. J Biomech Eng [Internet]. 2017 Mar 1 [cited 2020 Jan 21];139(3). Available from: http://www.ncbi.nlm.nih.gov/pubmed/27987301

9. Ploeger HE, Waterval NFJJ, Nollet F, Bus SA, Brehm MAA. Stiffness modification of two ankle-foot orthosis types to optimize gait in individuals with non-spastic calf muscle weakness-A proof-of-concept study. J Foot Ankle Res [Internet]. 2019 Aug 7 [cited 2020 Jan 21];12(1):41. Available from: http://www.ncbi.nlm.nih.gov/pubmed/31406508

10. Waterval NFJ, Nollet F, Harlaar J, Brehm MAA. Modifying ankle foot orthosis stiffness in patients with calf muscle weakness: Gait responses on group and individual level. J Neuroeng Rehabil [Internet]. 2019 Oct 17 [cited 2020 Jan 15];16(1):120. Available from: http://www.ncbi.nlm.nih.gov/pubmed/31623670

11. Sreenivasa M, Millard M, Felis M, Mombaur K, Wolf SI. Optimal control based stiffness identification of an ankle-foot orthosis using a predictive walking model. Front Comput Neurosci. 2017 Apr 13;11(April).

12. Alexander RM. Optimization and gaits in the locomotion of vertebrates. Physiol Rev [Internet]. 1989 [cited 2021 Dec 15];69(4):1199–227. Available from: https://pubmed.ncbi.nlm.nih.gov/2678167/

13. Bertram JEA, Ruina A. Multiple walking speed-frequency relations are predicted by constrained optimization. J Theor Biol [Internet]. 2001 Apr 21 [cited 2022 Feb 8];209(4):445–53. Available from: https://linkinghub.elsevier.com/retrieve/pii/S0022519301922799

14. Selinger JC, O’Connor SM, Wong JD, Donelan JM. Humans Can Continuously Optimize Energetic Cost during Walking. Curr Biol [Internet]. 2015 Sep 21 [cited 2022 Jan 23];25(18):2452–6. Available from: http://www.cell.com/article/S0960982215009586/fulltext

15. Kobayashi T, Orendurff MS, Hunt G, Lincoln LS, Gao F, LeCursi N, et al. An articulated ankle-foot orthosis with adjustable plantarflexion resistance, dorsiflexion resistance and alignment: A pilot study on mechanical properties and effects on stroke hemiparetic gait. Med Eng Phys [Internet]. 2017 Jun 1 [cited 2020 Jan 21];44:94–101. Available from: http://www.ncbi.nlm.nih.gov/pubmed/28284572

16. Bregman DJ. The Optimal Ankle Foot Orthosis. VU Amsterdam; 2011.

17. Arch ES, Stanhope SJ, Higginson JS. Passive-dynamic ankle-foot orthosis replicates soleus but not gastrocnemius muscle function during stance in gait: Insights for orthosis prescription. Prosthet Orthot Int [Internet]. 2016 Oct 1 [cited 2020 Jan 15];40(5):606–16. Available from: http://journals.sagepub.com/doi/10.1177/0309364615592693

18. Harper NG, Esposito ER, Wilken JM, Neptune RR. The influence of ankle-foot orthosis stiffness on walking performance in individuals with lower-limb impairments. Clin Biomech [Internet]. 2014 Sep [cited 2020 Jan 21];29(8):877–84. Available from: https://linkinghub.elsevier.com/retrieve/pii/S0268003314001739

19. Bregman DJJ, Van Der Krogt MMM, De Groot V, Harlaar J, Wisse M, Collins SHH. The effect of ankle foot orthosis stiffness on the energy cost of walking: A simulation study. Clin Biomech [Internet]. 2011 Nov [cited 2020 Jan 15];26(9):955–61. Available from: http://www.ncbi.nlm.nih.gov/pubmed/21723012

20. Waterval NFJ, Veerkamp K, Geijtenbeek T, Harlaar J, Nollet F, Brehm MA, et al. Validation of forward simulations to predict the effects of bilateral plantarflexor weakness on gait. Gait Posture [Internet]. 2021 Jun;87(December 2020):33–42. Available from: https://doi.org/10.1016/j.gaitpost.2021.04.020

21. Delp SL, Anderson FC, Arnold AS, Loan P, Habib A, John CT, et al. OpenSim: Open-source software to create and analyze dynamic simulations of movement. IEEE Trans Biomed Eng [Internet]. 2007 [cited 2020 Nov 29];54(11):1940–50. Available from: http://ieeexplore.ieee.org.

22. Seth A, Hicks JL, Uchida TK, Habib A, Dembia CL, Dunne JJ, et al. OpenSim: Simulating musculoskeletal dynamics and neuromuscular control to study human and animal movement. PLoS Comput Biol [Internet]. 2018 [cited 2020 Jan 13];14(7). Available from: https://doi.org/10.1371/journal.pcbi.1006223.g001

23. Geijtenbeek T. SCONE: Open Source Software for Predictive Simulation of Biological Motion. J Open Source Softw [Internet]. 2019 Jun 14 [cited 2020 Nov 22];4(38):1421. Available from: http://joss.theoj.org/papers/10.21105/joss.01421

24. Ong CF, Geijtenbeek T, Hicks JL, Delp SL. Predicting gait adaptations due to ankle plantarflexor muscle weakness and contracture using physics-based musculoskeletal simulations. PLoS Comput Biol [Internet]. 2019 Oct 1 [cited 2020 Jan 15];15(10):e1006993. Available from: https://doi.org/10.1101/597294

25. Delp SL, Loan JP, Hoy MG, Zajac FE, Topp EL, Rosen JM. An Interactive Graphics-Based Model of the Lower Extremity to Study Orthopaedic Surgical Procedures. IEEE Trans Biomed Eng. 1990;37(8):757–67.

26. Schillings ML, Kalkman JS, Janssen HMHA, van Engelen BGM, Bleijenberg G, Zwarts MJ. Experienced and physiological fatigue in neuromuscular disorders. Clin Neurophysiol. 2007;118(2):292–300.

27. Potvin JR, Fuglevand AJ. A motor unit-based model of muscle fatigue. PLoS Comput Biol. 2017 Jun 1;13(6).

28. Bigland-Ritchie B, Cafarelli E, Vollestad NK. Fatigue of submaximal static contractions. Acta Physiol Scand. 1986;128(SUPPL. 556):137–48.

29. Umberger BR, Gerritsen KGM, Martin PE. A model of human muscle energy expenditure. Comput Methods Biomech Biomed Engin. 2003;6(2):99–111.

30. Johnson MA, Polgar J, Weightman D, Appleton D. Data on the distribution of fibre types in thirty-six human muscles. An autopsy study. J Neurol Sci [Internet]. 1973 [cited 2021 Jan 4];18(1):111–29. Available from: https://pubmed-ncbi-nlm-nih-gov.tudelft.idm.oclc.org/4120482/

31. Garrett WE, Califf JC, Bassett FH. Histochemical correlates of hamstring injuries. Am J Sports Med [Internet]. 1984 Mar 23 [cited 2021 Jan 4];12(2):98–103. Available from: http://journals.sagepub.com/doi/10.1177/036354658401200202

32. Sherman MA, Seth A, Delp SL. Simbody: Multibody dynamics for biomedical research. In: Procedia IUTAM. Elsevier; 2011. p. 241–61.

33. Veerkamp K, Waterval NFJFJ, Geijtenbeek T, Carty CPP, Lloyd DGG, Harlaar J, et al. Evaluating cost function criteria in predicting healthy gait. J Biomech [Internet]. 2021 [cited 2021 Sep 13];123:110530. Available from: https://reader.elsevier.com/reader/sd/pii/S0021929021003110?token=C5F90B52ECE9BBADC350F4754674BC8D539DDEF6EA369F1101D492DDC8A5222D5F497C81239F5468BED23893568D4DE8&originRegion=eu-west-1&originCreation=20210913113928

34. Ries AJ, Schwartz MH. Ground reaction and solid ankle–foot orthoses are equivalent for the correction of crouch gait in children with cerebral palsy. Dev Med Child Neurol [Internet]. 2019 Feb 1 [cited 2021 Jan 12];61(2):219–25. Available from: http://doi.wiley.com/10.1111/dmcn.13999

35. Geyer H, Herr H. A Muscle-reflex model that encodes principles of legged mechanics produces human walking dynamics and muscle activities. IEEE Trans Neural Syst Rehabil Eng. 2010;18(3):263–73.

36. Hansen N. The CMA Evolution Strategy: A Comparing Review. In: Towards a New Evolutionary Computation [Internet]. Springer Berlin Heidelberg; 2007 [cited 2021 Jan 4]. p. 75–102. Available from: https://www.springerlink.com

37. Song S, Geyer H. A neural circuitry that emphasizes spinal feedback generates diverse behaviours of human locomotion. J Physiol [Internet]. 2015 Aug 15 [cited 2020 Nov 23];593(16):3493–511. Available from: http://doi.wiley.com/10.1113/JP270228

38. Uchida TK, Hicks JL, Dembia CL, Delp SL. Stretching your energetic budget: How tendon compliance affects the metabolic cost of running. Zadpoor AA, editor. PLoS One [Internet]. 2016 Mar 1 [cited 2020 Nov 18];11(3):e0150378. Available from: https://dx.plos.org/10.1371/journal.pone.0150378

39. Bril B, Ledebt A. Head coordination as a means to assist sensory integration in learning to walk. Neurosci Biobehav Rev [Internet]. 1998 Mar 4 [cited 2021 Jan 4];22(4):555–63. Available from: https://pubmed-ncbi-nlm-nih-gov.tudelft.idm.oclc.org/9595569/

40. Pozzo T, Berthoz A, Lefort L. Head stabilization during various locomotor tasks in humans. Exp Brain Res [Internet]. 1990 Aug [cited 2017 May 7];82(1):97–106. Available from: http://link.springer.com/10.1007/BF00230842

41. Ong CF, Hicks JL, Delp SL. Simulation-Based Design for Wearable Robotic Systems: An Optimization Framework for Enhancing a Standing Long Jump. IEEE Trans Biomed Eng [Internet]. 2016 May;63(5):894–903. Available from: http://ieeexplore.ieee.org/document/7173005/

42. Vicon Motion Systems. About the Plug-in Gait model - Nexus 2.9 Documentation - Vicon Documentation [Internet]. 2019 [cited 2022 May 17]. Available from: https://docs.vicon.com/display/Nexus212/About+the+Plug-in+Gait+model#AboutthePluginGaitmodel-PiGRefs

43. Whittle MW. Gait Analysis [Internet]. 4th ed. Gait Analysis. Elsevier; 2007 [cited 2021 Jan 5]. Available from: https://linkinghub.elsevier.com/retrieve/pii/B9780750688833X50016

44. MathWorks. Polynomial curve fitting - MATLAB polyfit - MathWorks United Kingdom [Internet]. MATLAB Documentation. 2016 [cited 2021 Jan 27]. Available from: https://nl.mathworks.com/help/matlab/ref/polyfit.html

45. Falisse A, Serrancolí G, Dembia CL, Gillis J, Jonkers I, De Groote F. Rapid predictive simulations with complex musculoskeletal models suggest that diverse healthy and pathological human gaits can emerge from similar control strategies. J R Soc Interface [Internet]. 2019 Aug 21 [cited 2020 Nov 22];16(157):20190402. Available from: https://royalsocietypublishing.org/doi/10.1098/rsif.2019.0402

46. Waters RL, Mulroy S. The energy expenditure of normal and pathologic gait. Gait Posture [Internet]. 1999 Jul 1 [cited 2021 Jan 20];9(3):207–31. Available from: https://linkinghub.elsevier.com/retrieve/pii/S0966636299000090

47. Brehm MA, Beelen A, Doorenbosch CAM, Harlaar J, Nollet F. Effect of carbon-composite knee-ankle-foot orthoses on walking efficiency and gait in former polio patients. J Rehabil Med. 2007 Oct;39(8):651–7.

48. Brehm MA, Harlaar J, Schwartz M. Effect of ankle-foot orthoses on walking efficiency and gait in children with cerebral palsy. J Rehabil Med. 2008 Jul;40(7):529–34.

49. Crowninshield RD, Brand RA. A physiologically based criterion of muscle force prediction in locomotion. J Biomech. 1981 Jan 1;14(11):793–801.

50. Ackermann M, van den Bogert AJ. Optimality principles for model-based prediction of human gait. J Biomech [Internet]. 2010 Apr [cited 2022 Feb 8];43(6):1055–60. Available from: http://dx.doi.org/10.1016/j.jbiomech.2009.12.012

51. Cook TM, Farrell KP, Carey IA, Gibbs JM, Wiger GE. Effects of restricted knee flexion and walking speed on the vertical ground reaction force during gait. J Orthop Sports Phys Ther [Internet]. 1997 [cited 2021 Jan 21];25(4):236–44. Available from: https://www.jospt.org

52. Zadpoor AA, Nikooyan AA. The relationship between lower-extremity stress fractures and the ground reaction force: A systematic review. Vol. 26, Clinical Biomechanics. Elsevier; 2011. p. 23–8.

53. Collins SH, Wiggin MB, Sawicki GS, Bruce Wiggin M, Sawicki GS. Reducing the energy cost of human walking using an unpowered exoskeleton. Nature [Internet]. 2015 Jun 1 [cited 2020 Jan 15];522(7555):212–5. Available from: http://www.nature.com/articles/nature14288

54. Matsubara JH, Wu M, Gordon KE. Metabolic cost of lateral stabilization during walking in people with incomplete spinal cord injury. Gait Posture. 2015;41(2):646–51.

55. Meyns P, Kerkum YL, Brehm MA, Becher JG, Buizer AI, Harlaar J. Ankle foot orthoses in cerebral palsy: Effects of ankle stiffness on trunk kinematics, gait stability and energy cost of walking. Eur J Paediatr Neurol [Internet]. 2020 [cited 2021 Feb 2];26:68–74. Available from: https://doi.org/10.1016/j.ejpn.2020.02.009

56. Milner CE, Ferber R, Pollard CD, Hamill J, Davis IS. Biomechanical factors associated with tibial stress fracture in female runners. Med Sci Sports Exerc [Internet]. 2006 Feb [cited 2021 Jan 27];38(2):323–8. Available from: http://journals.lww.com/00005768-200602000-00019

